# Improved production of SARS-CoV-2 spike receptor-binding domain (RBD) for serology assays

**DOI:** 10.1101/2020.11.18.388868

**Authors:** Jennifer Mehalko, Matthew Drew, Kelly Snead, John-Paul Denson, Vanessa Wall, Troy Taylor, Kaitlyn Sadtler, Simon Messing, William Gillette, Dominic Esposito

## Abstract

The receptor-binding domain (RBD) of the SARS-CoV-2 spike protein is a commonly used antigen for serology assays critical to determining the extent of SARS-CoV-2 exposure in the population. Different versions of the RBD protein have been developed and utilized in assays, with higher sensitivity attributed to particular forms of the protein. To improve the yield of these high-sensitivity forms of RBD and support the increased demand for this antigen in serology assays, we investigated several protein expression variables including DNA elements such as promoters and signal peptides, cell culture expression parameters, and purification processes. Through this investigation, we developed a simplified and robust purification strategy that consistently resulted in high levels of the high-sensitivity form of RBD and demonstrated that a carboxyterminal tag is responsible for the increased sensitivity in the ELISA. These improved reagents and processes produce high-quality proteins which are functional in serology assays and can be used to investigate seropositivity to SARS-CoV-2 infection.

Highlights:

- Improved yields of SARS-CoV-2 spike RBD through modification of DNA constructs and purification parameters
- Two versions of RBD show different sensitivity in serology assays
- Yields of greater than 50 mg/l obtained under optimal conditions
- Magnetic bead purification technology improves throughput of protein production

## Introduction

Serology assays are critical tools in the response to the COVID-19 pandemic [1]. Such assays can measure the presence and extent of an immune response and help to identify the number of asymptomatic SARS-CoV-2 infections in the population. Until an effective vaccine is developed, these are critical tools in identifying and controlling the spread of the infection. A number of serology assays have been published to date, with many employing subdomains of the SARS-CoV-2 S protein (hereafter referred to as spike) in ELISA-based assays. The specificity of the spike protein [1,2] makes it a clear target for therapeutic interventions such as vaccines or monoclonal antibodies, and also for use in serology studies to assess the prevalence of immune responses to the virus and therapeutics. In addition to soluble spike trimer, receptor binding domain (RBD) is also frequently utilized in these assays [3,4]. RBD, which interacts with the extracellular ACE2 receptor and permits entry of SARS-CoV-2 into cells, is considerably smaller and more readily generated in recombinant form than the full-length spike. While modified production methods have improved the production of soluble full-length spike protein [5], RBD production optimization has lagged. Recently, during development of assays to support an NIH-led serosurvey, it was shown that certain forms of RBD resulted in higher sensitivity ELISA results [6], presumably due to higher antibody affinity. To explore this further, and to optimize production of various RBD reagents, we investigated multiple protein constructs for protein production yield and their impact on the sensitivity of serology assays. This work allowed us to improve the production yield of the most sensitive form of RBD by modifying DNA sequence of the expression vectors, cell culture temperature and harvest time, and purification methodology. The final proteins produced were highly pure and functioned as sensitive and specific antigens in ELISAs. Our data also shows that improvements in serology assay sensitivity are caused the addition of a C-terminal streptavidin-binding protein (SBP) tag [7], which likely helps orient the protein on ELISA plates for better antibody detection. Taken together, these improvements allowed the production of sufficient RBD antigen for more than 6,000 ELISA plates per liter of culture.

## Materials and methods

### DNA

Original DNA for the expression of Sinai RBD [3] was generously provided by Dr. Florian Krammer (Icahn School of Medicine, Mt. Sinai) through BEI Resources, and is referred to here as construct X22. Original DNA for the expression of Ragon RBD [4] was generously provided by Dr. Aaron Schmidt (Ragon Institute of MGH, MIT, and Harvard), and is referred to here as construct X24. Modified DNA for both forms of RBD protein were generated by synthesis of Gateway Entry clones with gene optimization for mammalian expression (ATUM, Inc.). Entry clones were subcloned using Multisite Gateway recombination (ThermoFisher) into pDest-303 (Addgene #159678) with an optimized CMV51 promoter [8]. Final expression clones were validated by restriction analysis. The similarities and differences in these constructs are outlined in **Table 1** and shown schematically in **Fig. 1**. Transfection-quality DNA for all constructs was produced in-house using the Qiagen Plasmid Plus Maxi Kit per the manufacturer’s protocols or was generated at large-scale by Aldevron (Fargo, ND).

**Figure 1.**
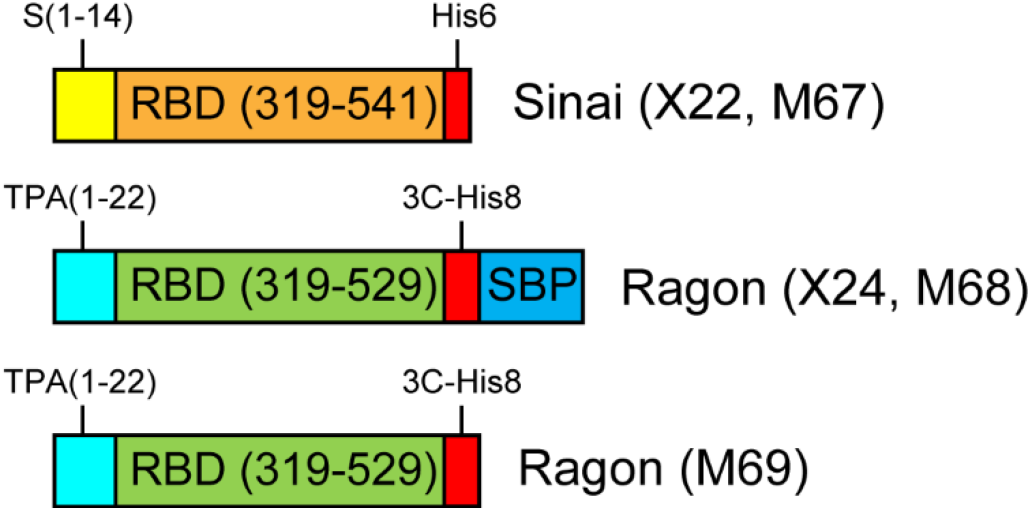
Comparison of RBD expression constructs. Four different gene designs were utilized in this work. All constructs contain the CoV-2 spike RBD domain, either containing amino acids 319-541 (Sinai, orange) or 319-529 (Ragon, green). All proteins contain a signal peptide for secretion of the RBD from mammalian cells—these leaders are from the CoV-2 spike protein (S, yellow) or human tissue plasminogen activator (TPA, cyan). Carboxyterminal tags are present on all constructs consisting of a 6 or 8 polyhistidine tag (with or without an HRV-3C protease cleavage site, red) and a streptavidin-binding protein tag (SBP, blue).

**Table 1.** Receptor-binding domain (RBD) constructs used in this work. Listed are the construct names/reference numbers, the amino acid region of SARS-CoV-2 spike utilized in the construct, the signal peptide (S = SARS-CoV-2 spike, TPA = human tissue plasminogen activator), the C-terminal tag attached to the RBD region (His6/His8 = polyhistidine tags, 3C = HRV3C protease cleavage site, SBP = streptavidin-binding peptide), the promoter used to drive transcription of the gene of interest (CAG = chicken beta-actin promoter with human CMV enhancer, HTLV = HTLV1 5’ UTR, CMV51 = enhanced cytomegalovirus immediate early promoter/enhancer), and the backbone vector utilized.

| Construct | Spike region | Sig peptide | C-tag | Promoter | Vector |
| --- | --- | --- | --- | --- | --- |
| X22 (Sinai) | 319-541 | S (1-14) | His6 | CAG | pCAGGS |
| M67 (FNL-S) | 319-541 | S (1-14) | His6 | CMV51 | pDest-303 |
| X24 (Ragon) | 319-529 | TPA (1-22) | 3C-His8-SBP | CMV-HTLV | pVRC |
| M68 (FNL-R) | 319-529 | TPA (1-22) | 3C-His8-SBP | CMV51 | pDest-303 |
| M69 (FNL-R) | 319-529 | TPA (1-22) | 3C-His8 | CMV51 | pDest-303 |

### Mammalian cell culture

Manufacturer’s protocols were followed for the transfection and culturing of Expi293F cells (Thermo Fisher Scientific, Waltham MA). Briefly, 1.7 liters of cell culture at 2.9 x 10^6^ cells/ml were transfected with preformed Expifectamine:DNA complexes at 1 μg/ml of final culture volume. Expression cultures were incubated at 37°C and 8% CO_2_ in 5-liter Optimum Growth Flasks (Thomson Instrument Company, Oceanside, CA) shaking at 105 RPM on an Infors HT Multitron Standard orbital shaker with a 2” orbit. Expression enhancers were added 18-20 hours post-transfection per manufacturer’s instructions, and incubation was continued at 37°C and 8% CO_2_ until harvest time of either 72, 96, or 120 hr post-transfection. For temperature shift experiments, the incubation temperature was lowered to 32°C immediately after enhancer addition.

### Tangential flow filtration (TFF)

Harvested culture supernatants were clarified by centrifugation (4000 x *g*, 20 min, 4°C) followed by filtration (catalog# 12993, Pall Corporation, Port Washington, NY). When used in column chromatography, clarified supernatants were concentrated and buffer exchanged by TFF. Specifically, a MasterFlex peristaltic pump (Vernon Hills, IL) fed the clarified supernatant to a 10 kDa MWCO cassette (catalog# SK1P003W4, MilliporeSigma, Burlington, MA). The clarified supernatant was concentrated to 10% of the initial volume, and then buffer exchanged with 5 volumes of 1x PBS, pH 7.4 (Buffer A, diluted from 10X PBS, catalog #70011069 Thermo Fisher Scientific, Waltham, MA). Following buffer exchange, the TFF cassette was rinsed with 200-250 ml Buffer A, to collect any protein remaining in the cassette. Clarified supernatants for use in batch purification were not buffer exchanged.

### Standard protein purification

Chromatography was conducted at room temperature (~22°C) using NGC medium-pressure chromatography systems from BioRad Laboratories Inc. (Hercules, CA) with exceptions noted below. The standard purification protocol employed immobilized metal affinity chromatography (IMAC) and size exclusion chromatography (SEC). Specifically, TFF-treated culture supernatant was adjusted to 25 mM imidazole and applied to a 10 ml Ni Sepharose High Performance nickel-charged column (GE Healthcare, Chicago, IL) previously equilibrated in Buffer A + 25 mM imidazole. The flow rate for all steps of the IMAC was 5 ml/min. The column was washed in Buffer A + 25 mM imidazole for 4 column volumes (CV) with the final 3 CV collected separately as the column wash. The protein was eluted from the column by applying a 20 CV linear gradient of 25 mM – 500 mM imidazole in Buffer A followed by a 3 CV step elution of Buffer A + 500 mM imidazole. Fractions (5 ml) were collected for all elution steps. Elution fractions were analyzed by SDS-PAGE/Coomassie-staining and appropriate fractions were pooled. Typical pool volume for a purification from one liter of culture supernatant was ~80 ml.

The sample was concentrated using Amicon Ultra Spin Concentrators with a 10 kDa molecular weight cut off membrane (Millipore, MA, USA), discarding the permeate. Once retentate volume has reached 5 ml, the protein concentration was determined by measuring the A280 using a Nanodrop One spectrophotometer (Thermo Fisher Scientific, MA, USA). This protein was applied to a 16/600 Superdex 75 preparative size exclusion column (GE Healthcare, Chicago, IL) previously equilibrated in Buffer A. The flow rate was 1 ml/min, and the protein was eluted with 1 CV of Buffer A. 1 mL fractions were collected starting at 0.2 CV of elution. Elution fractions were then analyzed by SDS-PAGE/Coomassie-staining and appropriate monomeric fractions were pooled. Typical pool volume from one liter of starting material was ~15 ml. The protein concentration was determined by measuring the A280 using a Nanodrop One spectrophotometer, and final pool was filtered through a 0.22 μm syringe filter with low protein binding capacity. Final protein was snap frozen in liquid nitrogen in 0.5 ml aliquots, and stored at −80°C.

For comparing Ragon protein with and without the SBP fusion tag, a 20 mg sample of the X24 protein was removed after the initial IMAC chromatography step and incubated with 2 mg rhinovirus 3C protease overnight at 25°C, while dialyzing to 1x PBS, pH 7.4. The next day, this sample was subjected to subtractive IMAC washing with 1x PBS, pH 7.4 containing 25 mM imidazole. Cleaved X24 eluted in the flowthrough fractions and was pooled and subjected to SEC under the same conditions as the full-length X24 protein above.

For SDS-PAGE analysis of purified proteins, 20 ug of each purified sample was brought to a final volume of 20 ul in water. 4 ul of PNGaseF buffer was then added, along with 1 ul PNGaseF (ThermoFisher), for a final volume of 25 ul. Samples were incubated at 50°C for 5 min and 5 ul of each were electrophoresed using SDS-PAGE/Coomassie-staining and compared with proteins which were untreated.

### Magnetic Bead protein purification

For the batch purification from filtered (see above) culture supernatants, 0.5 ml of Ni-charged MagBeads (GeneScript, Piscataway, NJ), previously equilibrated in Buffer A, were placed in the bottom of each of two 50 ml conical tubes and filtered culture medium was added to the tubes (40 ml per tube). Tubes were incubated at room temperature on an orbital mixer for 1 hr. After incubation, a rare-earth magnet was used to capture the beads to the side of the conical tubes, and the medium was removed and saved as “flow through”. Beads were then combined into one 50 ml conical tube using 5 ml Buffer A, and then 10 ml of Buffer A was added. This first wash was incubated on the orbital mixer at room temperature for 5 min. Beads were collected, the wash removed, and a second 15 ml wash step was performed. Beads were then transferred to a 5 ml snap-cap MacroTube (MTC Bio) for elution steps using 3 ml Buffer A. The buffer was then decanted. Protein was eluted from the washed beads by addition of 2 ml Buffer A + 500 mM imidazole. Each elution was incubated on the orbital mixer for 30 minutes at room temperature, and then collected in a 15 ml conical tube. This process was repeated for a total of 3 elutions.

Samples of each elution fraction were analyzed by SDS-PAGE/Coomassie-staining and appropriate fractions were pooled. Typically, all 3 elutions were pooled for a 6 ml total elution volume. The protein concentration was determined by measuring the A280 using a Nanodrop One spectrophotometer, and final protein was snap frozen in liquid nitrogen, and stored at −80°C.

### Enzyme-linked immunosorbent assay (ELISA)

In order to assess antigen sensitivity, purified RBD proteins (including cleaved Ragon X24 lacking the SBP tag) were used as antigens in an ELISA with positive control SARS-CoV-2 monoclonal antibody to the RBD domain. The ELISA was carried out as previously reported [6] using 200 ng per well of the various RBDs and serial 5-fold dilutions of a 2.5 ug/ml stock of SARS-CoV RBD monoclonal antibody mAb109 (construct kindly provided by the NIAID Vaccine Research Center).

## Results and Discussion

To produce SARS-CoV-2 RBD for the development of serology assays, we initially tested two constructs previously published in the literature [3,4]. Initial findings showed that the construct from the Ragon Institute, although containing less spike protein sequence, resulted in more than a 2-fold higher sensitivity in ELISA assays than the Mt. Sinai version of the protein [6]. However, the Ragon protein production yield was considerably lower than that of the Mt. Sinai protein, even though the Ragon protein contained an additional tag sequence which should have produced, by mass, more protein. This suggested to us that there were suboptimal features of the expression system, which could be optimized for the Ragon protein. We also were interested in determining why the sensitivity of the Ragon construct was higher in the ELISA assays, and whether this was a result of the different portion of spike RBD in these proteins or the added tag sequences. Therefore, we generated novel DNA expression constructs to optimize the purification process and to improve the production yield and quality of these vital protein reagents.

### DNA construct improvements

Comparison of the expression vectors used for production of the two RBD constructs showed that there were few similarities in the composition of the two DNAs, making direct comparison difficult. For this reason, we decided to reclone both proteins with the same C-terminal tags originally used, but in identical backbone vectors with the same promoter. To this end, we acquired optimized DNA constructs from ATUM, designed to maximize expression in HEK293 cells, and transferred those genes to our highly optimized mammalian expression vector pDest-303, with a strong CMV51 promoter which provides high-level protein production in HEK293 cells. This was done for both the Ragon protein (M68) and the Sinai protein (M67). In addition, to further explore possible roles for tagging, we engineered an additional version of the Ragon protein which removed the SBP tag (M69). These 5 constructs are highlighted in **Fig. 1**, and details about their construction are noted in **Table 1**.

### Optimization of protein expression and purification

Previously, we had optimized production of SARS-CoV-2 soluble spike protein using the Expi293 expression system with reduced temperature to enhance protein secretion [5]. A similar strategy was employed with RBD proteins, using a 32°C expression for 96 hours to maximize production while maintaining high levels of cell viability. We investigated whether longer incubation might also enhance protein yield, but harvest at 120 hours post-transfection did not improve production, and in some cases, may have actually reduced final yields (data not shown).

Published protocols for purification of RBD domains often utilize a batch process, which generally is less efficient than column chromatography. For this reason, we developed a modified approach to that used for soluble spike production, including an initial tangential flow filtration step to buffer exchange and concentrate the initial supernatant, followed by a column IMAC process with a gradient elution. Protein was subsequently concentrated and applied to a size exclusion column for final polishing. In general, all constructs performed similarly in this process regardless of expression level, and all protein was readily captured by the IMAC resin. A typical purification is shown in **Fig. 2A** (IMAC) and **Fig. 2B** (SEC) for the M68 protein.

**Figure 2.**
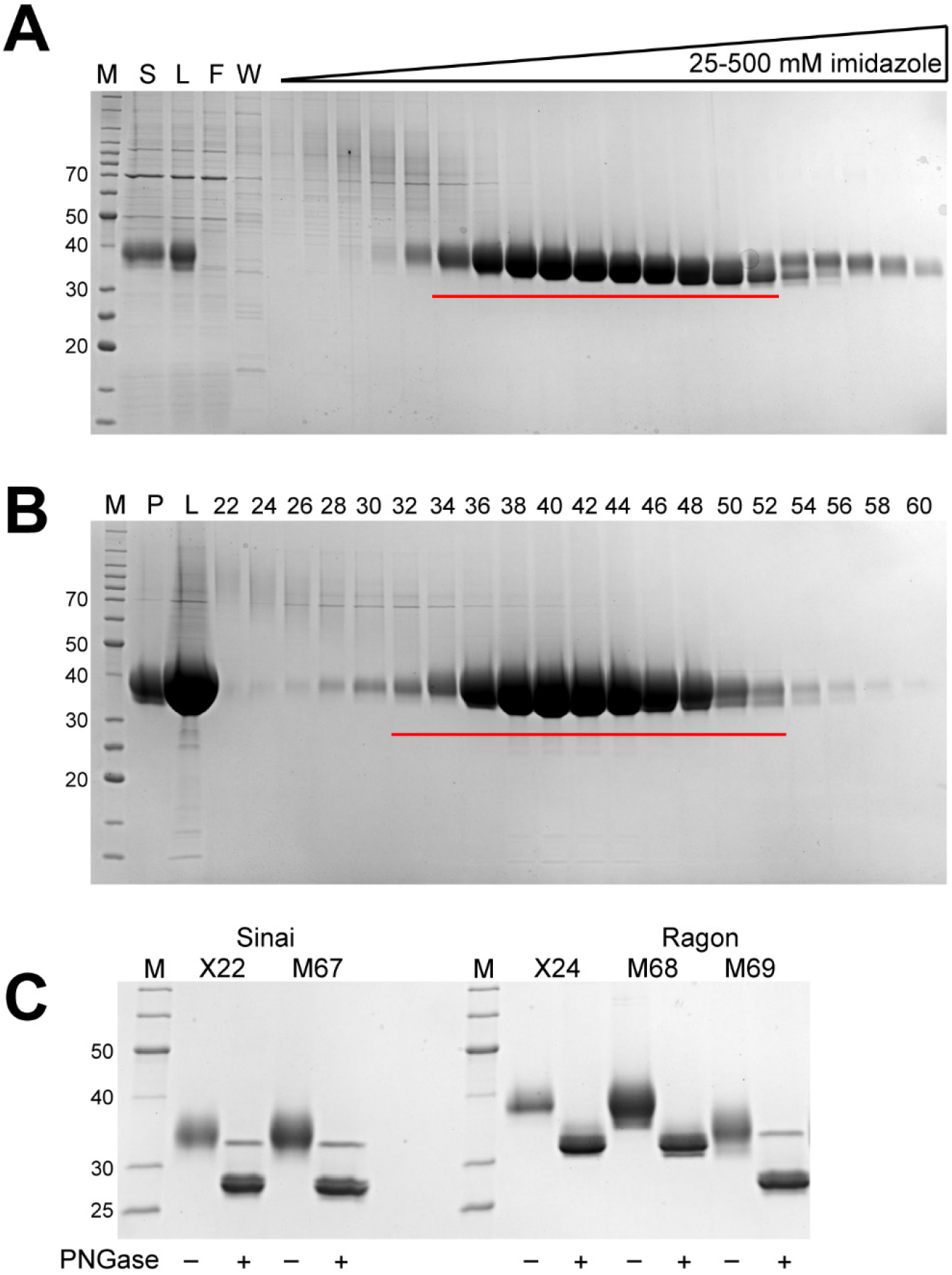
Representative chromatography fraction analyses and purified proteins. M – protein standards, molecular weights of select standards noted in kDa. **A.** Coomassie-stained SDS-PAGE analysis of fractions from IMAC chromatography of the Ragon M68 protein, S – culture supernatant, L – TFF retentate/column load, F – column flow through, W – column wash. Fractions pooled are underlined in red. **B.** SDS-PAGE/Coomassie staining analysis of fractions from SEC chromatography of the Ragon M68 protein, P – IMAC pool, L – concentrated pool/column load, numbers represent fraction numbers (every other fraction loaded). Fractions pooled for the final protein sample are underlined in red. **C.** SDS-PAGE/Coomassie staining analysis of final purified proteins. Alternating lanes are without or with the addition of PNGase F to remove N-linked glycans.

Final proteins from one complete set of purifications are shown in **Fig. 2C**, and a summary of purification yields from multiple independent productions of each protein are highlighted in **Table 2**. From this data, it is clear that yields from our optimized vector system are higher than either original construct. In the case of the Sinai protein, the yield improvement averaged 1.4x, while the Ragon protein levels were increased nearly 4x by these vector modifications. This data is consistent with our expectations based on the promoters used in the original vectors and our experience with pDest-303. Notably, the yields of the non-SBP containing M69 protein was similar to that of the M67 Sinai construct—all of these proteins are nearly identical in size and amino acid sequence and would be expected, under optimal conditions, to produce similar levels of protein. If the SBP-containing M68 is corrected for the larger mass of the fusion protein, the 63.6 mg/l yield is effectively the same molar productivity as the other constructs, again suggesting that these RBD proteins are being produced at similar levels regardless of construct details. **Fig. 2C** also highlights the glycosylation observed on the purified RBD proteins, as clearly noted by the smeared appearance of the purified proteins. After treatment with PNGase F to remove N-linked glycans known to be present in spike at Asn331 and Asn343, the proteins migrate at a more appropriate molecular weight on the SDS-PAGE gel and with a more defined appearance.

**Table 2.** Protein yields of various RBD constructs using the standard TFF-IMAC-SEC process. Numbers represent yields from independent experiments. RBDs represents the same RBD amino acid sequence as the Sinai construct, while RBDr represents the same RBD amino acid sequence as the Ragon construct.

| Condition | Protein Yield (mg/l) | Mean (mg/l) |
| --- | --- | --- |
| X22 (Sinai) | 29.6, 30.6, 40.9 | 33.7 ± 6.2 |
| M67 (RBDs-His6) | 45.1, 50.1 | 47.6 ± 3.5 |
| X24 (Ragon) | 8.1, 8.3, 8.6, 11.9, 12.2, 12.6, 18.9 | 11.5 ± 3.8 |
| M68 (RBD<sub>r</sub>-3C-His8-SBP) | 58.5, 58.7, 73.6 | 63.6 ± 8.7 |
| M69 (RBD<sub>r</sub>-3C-His8) | 45.0, 47.7, 55.7 | 49.5 ± 5.6 |

### Improved purification yield using IMAC magnetic beads

Our observations with full-length soluble spike protein [5] suggested that an alternative batch purification method utilizing magnetic bead technology could improve yields and significantly reduce purification time and cost. Thus, we used magnetic IMAC beads to capture RBD in batch mode from filtered lysates. A typical purification for M68 is shown in **Fig. 3A** where the majority of the protein was bound to the MagBeads and was eluted in the first two imidazole-containing fractions. The final proteins purified with this approach were similar in terms of quantity and final purity (**Table 3** and **Fig. 3B**) to proteins purified by the TFF/IMAC/SEC column process. This batch process has several distinct advantages over the more complex protocol including the elimination of the labor-intensive TFF process, which for large-scale transient cultures can take many hours, and reduced time and cost for column chromatography. The IMAC beads are also readily scaled across large volumes, making this a more consistent approach when culture conditions are varied. It is interesting to note that the original Sinai construct (X22) gives nearly double the protein yield using the MagBead process than expected from the standard purification. Since protein quality appears similar, this may suggest that a significant portion of the Sinai protein is being lost in the standard purification during the concentration or SEC steps which are not present in the MagBead production. Yields of the Ragon protein appear to be generally similar in the two processes.

**Table 3.** Protein yields of various RBD constructs using the MagBead process. Numbers represent yields from independent experiments. RBDs represents the same RBD amino acid sequence as the Sinai construct, while RBDr represents the same RBD amino acid sequence as the Ragon construct. n/a: not applicable as only one purification was performed.

| Condition | Protein Yield (mg/l) | Mean (mg/l) |
| --- | --- | --- |
| X22 (Sinai) | 62.1, 72.5 | $67.3 \pm 7.4$ |
| M67 (RBDs-His6) | 79.8 | n/a |
| X24 (Ragon) | 15.6, 19.2, 21.8 | $18.9 \pm 3.1$ |
| M68 (RBD<sub>r</sub>-3C-His8-SBP) | 78.3 | n/a |
| M69 (RBD<sub>r</sub>-3C-His8) | 58.8 | n/a |

**Figure 3.**
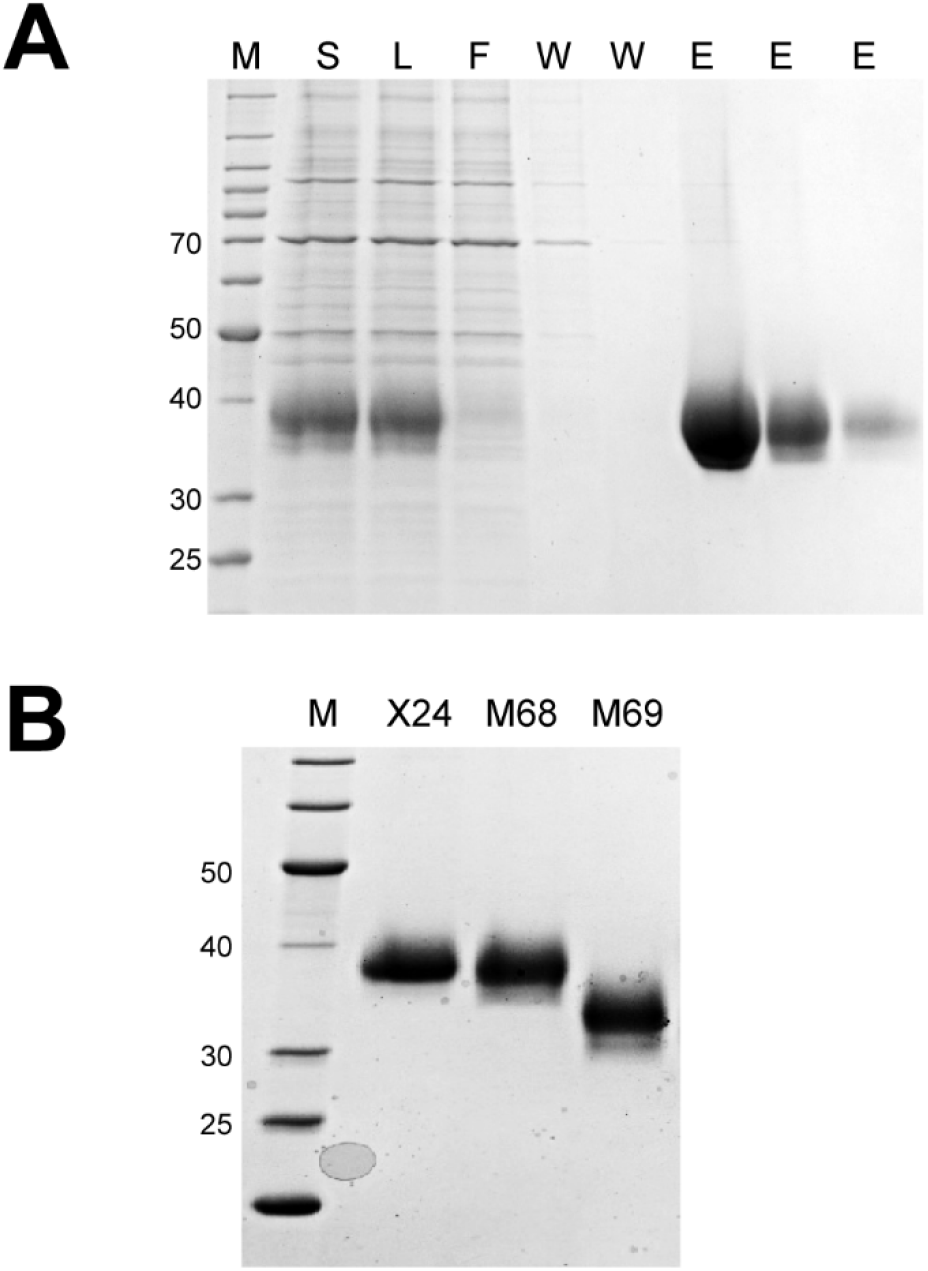
Representative magnetic bead chromatography analysis and purified proteins. M – protein standards, molecular weights of select standards noted in kDa. **A.** Representative SDS-PAGE/Coomassie staining analysis of IMAC magnetic bead chromatography of the Ragon X24 protein. S – culture supernatant, L – filtered culture supernatant/bead load, F – bead “flow through”, W – bead washes, E – bead elutions. **B.** SDS-PAGE/Coomassie staining analysis of MagBead purified protein pools.

### Performance of proteins in serology assays

We have previously developed and optimized a serology assay using an ELISA format [6], which showed that the Ragon version of RBD was more sensitive than the Sinai version. To identify what components of our newly produced RBD proteins might lead to this higher ELISA sensitivity, we compared all of the proteins in the serology assay. **Fig. 4** confirms that the Ragon proteins (X24, open squares; M68, filled squares) were more sensitive than the Sinai (X22) protein (open triangles)—in this case a 5-fold increase in antibody concentrations which give equivalent signals. It appears that the C-terminal SBP tag is entirely responsible for this higher sensitivity, as the cleaved Ragon X24 protein (open circles) and the Ragon M69 protein lacking the SBP tag (filled circles) both display similar low sensitivity as the Sinai RBD. The particular amino acid boundaries of the RBD domain do not seem to play a role in the difference in sensitivity based on our results. We propose that the SBP tag interacts with the ELISA plate to uniquely position the RBD domain to afford greater accessibility of antibodies to the RBD. It would be interesting to see if tags other than SBP might provide the same enhancement, or if there is a specific feature of this tag which uniquely results in greater sensitivity. In any case, we demonstrate here that the presence of the SBP tag on these constructs significantly improves sensitivity of the assay, making it the optimal choice for use in SARS-CoV-2 serology efforts.

**Figure 4.**
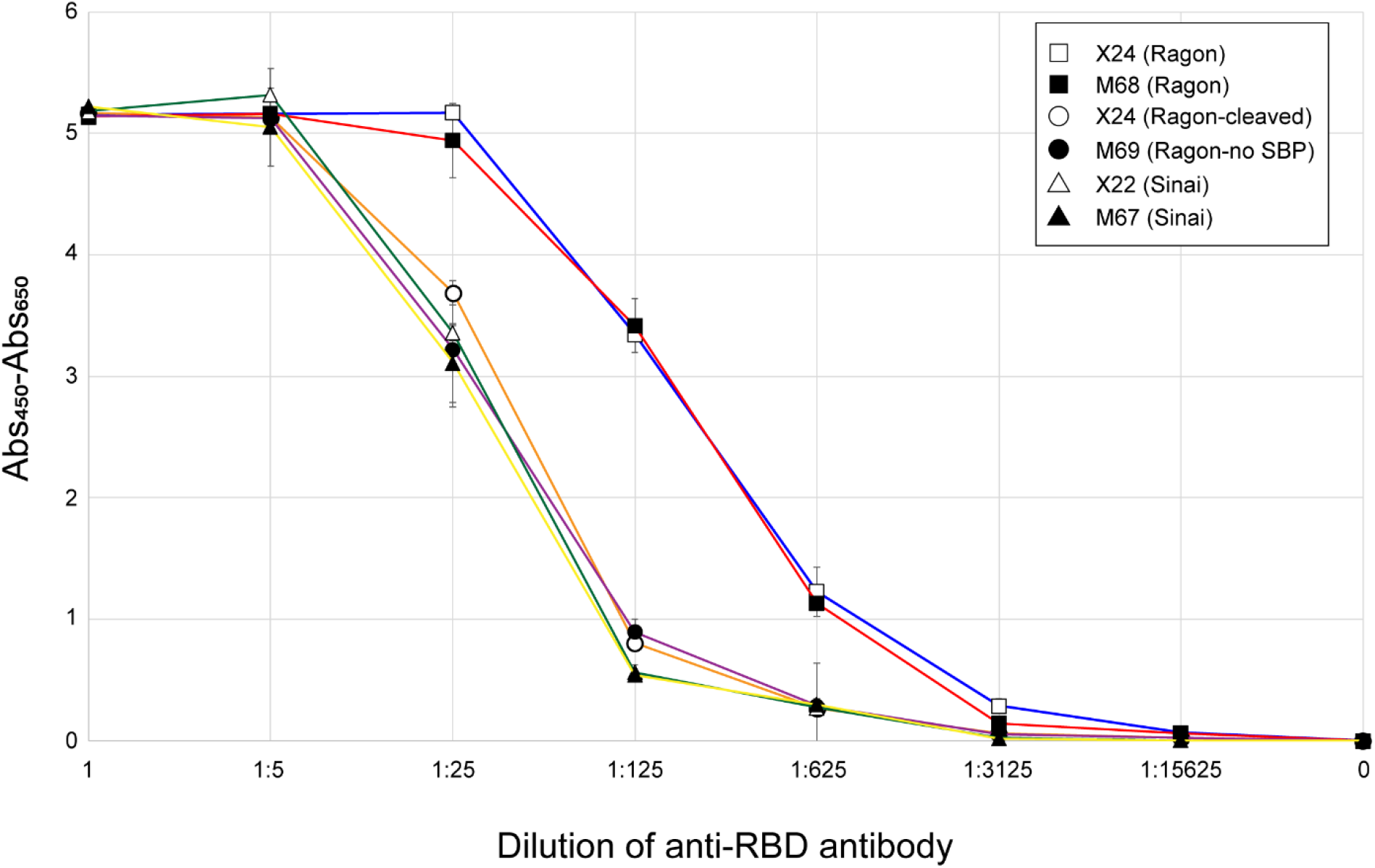
ELISA sensitivity of purified RBDs. The five RBD proteins, as well as a 3C protease-treated version of X24, were used to coat ELISA plates which were then treated with positive control anti-RBD monoclonal antibodies at the indicated dilutions. All measurements were performed in triplicate and means are plotted with standard deviations noted with error bars. Measurements are based on absorbance at 450 nm corrected by subtraction of absorbance at 650 nm. Samples tested were Sinai X22 (open triangles), Sinai M67 (filled triangles), Ragon X24 (open squares), Ragon X24 cleaved with 3C protease (open circles), Ragon M68 (filled squares), and Ragon M69 (filled circles).

We also compared the performance of X24 protein purified using the standard purification procedure (**Fig. 5**, filled squares and solid line) and the MagBead procedure (**Fig. 5**, open squares and dashed line) and show that both proteins perform equally in the ELISA. This demonstrates that the lower cost and more rapid MagBead purification process can be used to generate high-quality proteins for serology assays.

**Figure 5.**
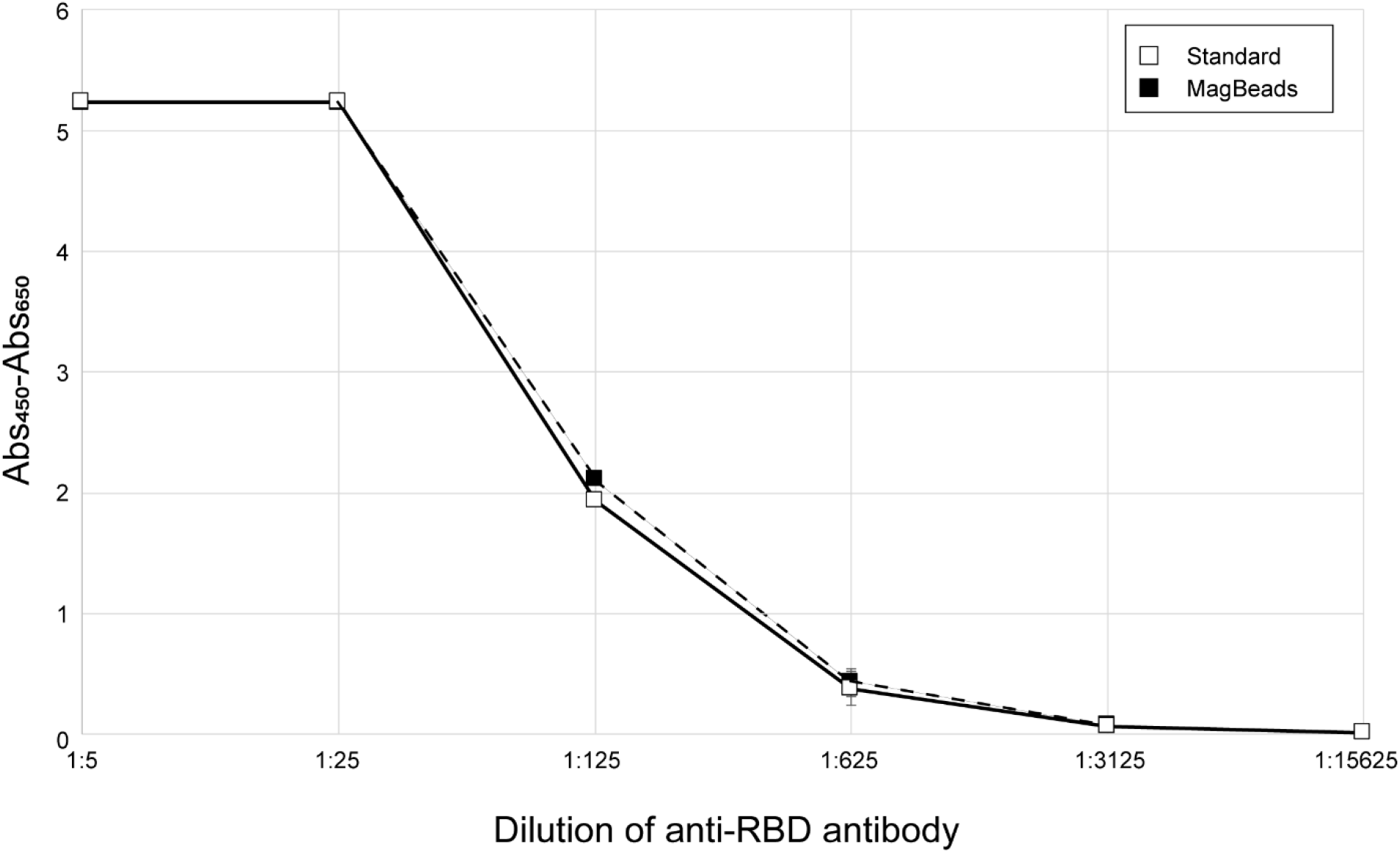
Comparison of *ELISA sensitivity of RBD purified by two different methods*. The Ragon X24 protein purified by the standard process (open squares, solid line) or the MagBead process (filled squares, dashed line) were used to coat ELISA plates which were then treated with positive control anti-RBD monoclonal antibodies at the indicated dilutions. All measurements were performed in triplicate and means are plotted with standard deviations noted with error bars. Measurements are based on absorbance at 450 nm corrected by subtraction of absorbance at 650 nm.

## Conclusions

In summary, we presented multiple improvements to the production of SARS-CoV-2 RBD that increase the ability of laboratories to generate this high-quality vital reagent for serology assays and other applications. In particular, we confirmed that the SBP tag present on the Ragon RBD construct makes it more attractive as a serology assay substrate due to the higher level of sensitivity. Additional work needs to be done to clarify the specific role of SBP in the process. Deployment of this reagent for serology assays to determine the prevalence of SARS-CoV-2 infection are currently underway.

## Acknowledgements

We acknowledge Dr. Matthew Hall (NIH/NCATS) and Dr. Matthew Memoli (NIH/NIAID) for their role in the development of the serology assays used to validate these proteins. The authors would like to acknowledge Dr. Aaron G. Schmidt, Jared Feldman, Blake M. Hauser and Timothy M. Carradonna at the Ragon Institute of MGH, MIT and Harvard for providing the original plasmids for protein expression. This research was supported in part by the Intramural Research Program of the National Institute for Biomedical Imaging and Bioengineering. This project has been funded in whole or in part with Federal funds from the National Cancer Institute, National Institutes of Health, under contract number HHSN261200800001E. The content of this publication does not necessarily reflect the views or policies of the Department of Health and Human Services, nor does mention of trade names, commercial products, or organizations imply endorsement by the U.S. Government.

## Author Contributions

Conceptualization: JM, MD, TT, KSa, SM, WG, DE; Formal analysis: JM, MD, DE; Funding acquisition: DE; Investigation: JM, MD, KSn, JD, VW, TT; Methodology: JM, MD, KSn, VW, TT, SM, WG, DE; Project administration: WG, DE; Resources: MD, KSa, WG, DE; Supervision: MD, SM, WG, DE; Validation: JM, MD, KSn, JD, VW, TT, SM, WG, DE; Visualization: JM, MD, WG, DE; Writing – original draft: JM, MD, WG, DE; Writing – review and editing: JM, MD, KSa, SM, WG, DE.

## Abbreviations

AnSEC: analytical size exclusion chromatography
CV: column volume
ELISA: enzyme-linked immunosorbent assay
IMAC: immobilized metal ion affinity chromatography
MWCO: molecular weight cut-off
RBD: receptor binding domain
SBP: streptavidin-binding peptide
SDS-PAGE: sodium dodecyl sulfate-polyacrylamide gel electrophoresis
SEC: size exclusion chromatography
TFF: tangential flow filtration

